# Association between maternal depressive symptoms and hair cortisol concentration during pregnancy with corpus callosum integrity in newborns

**DOI:** 10.1101/2024.09.06.610927

**Authors:** Isabella L.C Mariani Wigley, Paula Mustonen, Linnea Karlsson, Saara Nolvi, Noora M Scheinin, Susanna Kortesluoma, Massimiliano Pastore, Katja Tervahartiala, Bárbara Coimbra, Ana João Rodrigues, Nuno Sousa, Hasse Karlsson, Jetro J Tuulari

**Affiliations:** FinnBrain Birth Cohort Study, Turku Brain and Mind Center, Department of Clinical Medicine, University of Turku, Finland; Department of Child Psychiatry, University of Turku and Turku University Hospital, Turku, Finland; Department of Clinical Medicine, Department of Public Health, University of Turku and Turku University Hospital; Centre for Population Health Research, Turku University Hospital and University of Turku; Department of Psychiatry, University of Turku and Satakunta Wellbeing Services County, Turku, Finland; Department of Psychiatry, University of Turku and Turku University Hospital, Turku, Finland; Department of Developmental and Social Psychology, University of Padua, Padua, Italy; Department of Psychology and Speech-Language Pathology, University of Turku, Turku, Finland; Department of Psychology, University of Jyväskylä, Jyväskylä, Finland; Life and Health Sciences Research Institute (ICVS), School of Medicine, University of Minho, Braga, Portugal; Clinical Neurosciences, University of Turku, Turku, Finland; Neurocenter, Turku University Hospital, Turku, Finland; Clinical Academic Center (2CA), Braga, Portugal

**Author notes:** Correspondence: Mariani Wigley Isabella L.C < >. These authors contributed equally to this work (Isabella L.C Mariani Wigley, Paula Mustonen).

**Keywords:** maternal depressive symptoms, hair cortisol, DTI, corpus callosum, pregnancy, newborns

## Abstract

**Background:** Maternal prenatal depressive symptoms are linked to neurodevelopmental impairments in offspring. Maternal cortisol levels are hypothesized to moderate this association, but its relationship with depressive symptoms is inconsistent. This study examined how maternal prenatal depressive symptoms and cortisol levels predict infant brain development, focusing on neonatal corpus callosum (CC) integrity.

**Methods:** Using data from the FinnBrain Birth Cohort Study, we analyzed 37 mother-infant dyads. MRI data were collected from 2 to 5 weeks old infants, and DTI imaging estimated fractional anisotropy (FA) in CC regions (Genu, Body, and Splenium). Maternal cortisol levels were assessed through hair cortisol concentration (HCC) from a 5cm hair segment, reflecting cortisol over the last five months of pregnancy. A factor score of maternal depressive symptoms was computed from EPDS questionnaire data collected at gestational weeks 14, 24, and 34. We employed multivariate regression models with a Bayesian approach for statistical testing, controlling for maternal and infant attributes.

**Results:** Results indicated that maternal prenatal depressive symptoms and HCC interact negatively in predicting infants’ FA across all CC regions. Infants exposed to high prenatal depressive symptoms and low HCC (1 SD below the mean) showed higher FA in all CC regions.

**Conclusions:** These findings highlight the complex dynamics between maternal prenatal cortisol levels and depressive symptoms, revealing a nuanced impact of those factors on the structural integrity of infants’ CC.

## 1. Introduction

The intricate relationship between maternal well-being during pregnancy and fetal development has long fascinated researchers due to its significant implications for offspring health and development^1^. This area of study is deeply rooted in the principles of Developmental Origins of Health and Disease (DOHaD) and Fetal Programming, which suggest that prenatal environmental exposures can profoundly influence future health outcomes by affecting various physiological systems^2,3^. These theories propose that critical periods during fetal development are particularly susceptible to external influences, which can have lasting effects on health trajectories, including the risk of developing chronic diseases later in life. Among these influences, the brain —known for its complexity and plasticity— plays a central role, with particular attention given to its white matter (WM)^4^.

Diffusion tensor imaging (DTI) has become the primary technique for studying WM microstructure, especially in early life^5^. This imaging modality provides metrics such as fractional anisotropy (FA), which measures the directionality and coherence of water diffusion in brain tissue^6^. FA is sensitive to changes in WM microstructure, which is continuously shaped throughout life and thus remains susceptible to early-life influences^6^, with potential cascading effects on subsequent brain development and function. Among the various factors impacting WM development, maternal mental health has emerged as a crucial focus^7^. For instance, alterations in WM microstructure have been observed in offspring of mothers experiencing prenatal depression^8^. Maternal depressive symptoms can affect physiological stress responses, thereby disrupting the intrauterine environment and impacting both fetal and postnatal development^9^.

The prevalence of depressive symptoms during pregnancy ranges from 10% to 20% in Western countries^10^. Evidence links prenatal exposure to maternal depression with adverse neurobehavioral outcomes in children, such as emotional and behavioral difficulties, and alterations in brain structure and function^9^. Despite these associations, the mechanisms underlying these links remain largely unknown. Previous studies have investigated neural characteristics in infants born to mothers with depressive symptoms. Posner et al. (2016)^11^ observed reduced structural connectivity between the right amygdala and the right ventral prefrontal cortex in infants of mothers with elevated prenatal depressive symptoms. Dean et al. (2018) reported higher diffusivity in right frontal regions in one-month-old infants of mothers with greater prenatal depressive and anxious symptoms. Notably, they also found a positive relationship between maternal symptoms and FA values in the corpus callosum (CC) specifically in males^12^. The CC, a major WM structure, is crucial for interhemispheric communication and integrates sensory, motor, and cognitive information^13,14^.

The CC plays a significant role in connecting various brain regions and contributing to networks like the default mode and salience networks^13,15,16^, which are considered critical for the susceptibility to environment^17^. Structural changes in the CC have been linked to prenatal and early-life stressors, including maternal anxiety and depression during pregnancy^18–20^, as well as childhood maltreatment^21^. These changes have been also associated with various developmental outcomes, including high emotional reactivity in infants, and a range of psychiatric conditions in later life, such as depression and schizophrenia^22–27^. Alterations in CC microstructure, especially decreased FA, are also observed in adults with treatment-resistant depression^28^, underscoring the significance of CC microstructure in susceptibility to environmental influences throughout the lifespan. Given the established links between maternal depressive symptoms and infant CC integrity, and the potential for at-risk developmental outcomes, it is essential to understand the mechanisms underpinning these associations. Consequently, it becomes imperative to expand the vision and integrate other key factors into the research framework.

Cortisol, the end-product hormone of the hypothalamic-pituitary-adrenal (HPA) axis, is often considered a proxy for maternal psychological distress^29,30^. Measuring cortisol from hair samples has become a standard method for assessing long-term systemic cortisol exposure, providing insights into maternal stress physiology over extended periods^31^. Emerging research on maternal hair cortisol concentration (HCC) has revealed associations between perinatal HCC and patterns of neural activity in infants, as measured by EEG at 6-12 months^32^. These findings also show that increased maternal perinatal HCC is linked with microstructural alterations in the amygdala in boys and changes in structural connectivity in girls^33^. Despite these advances, research on the joint effects of maternal depressive symptoms and HCC on newborn WM microstructure remains limited and we can be confident enough to state that, thus far, none of these studies have investigated the effects of the interaction between maternal depressive symptoms and HCC on the microstructure of the newborn’s WM.

To address this gap, the present study aims to explore the interplay between maternal prenatal depressive symptoms, HCC, and infant WM. Given the role of the CC in previous studies linking maternal depressive symptoms with WM organization, our focus is on the Genu, Body, and Splenium of the CC (Figure 1). The reliable delineation of the CC in the developing brain enhances the methodological robustness of this research^5,34,35^. Our hypothesis is that maternal prenatal depressive symptoms and HCC will interact in predicting the development of the CC in offspring. Given the variability in previous studies and the absence of prior research directly addressing this interaction, we have chosen not to speculate on whether lower or higher FA values will reflect susceptibility to maternal depressive symptoms and cortisol levels that HCC proxies.

**Figure 1.**
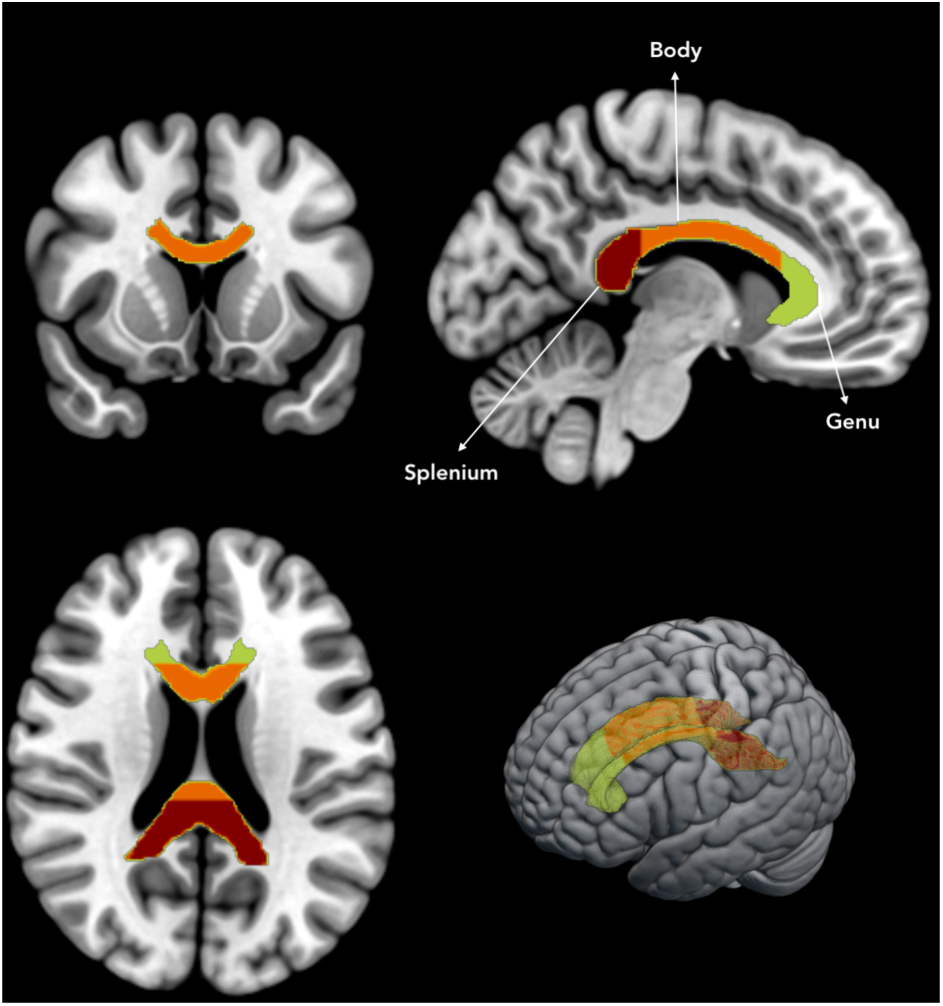
Corpus Callosum Sections. All figures are presented in radiological convention (left is right and right is left). Anatomical labels have been provided as per the JHU atlas. Colors represent corpus callosum genu (green), body (orange) and splenium (red).

## 2. Methods

### 2.1 Study population

This convenience sample is part of the FinnBrain Birth Cohort Study (www.finnbrain.fi;^36^). Recruitment occurred between December 2011 and April 2015 in Turku and the Åland Islands, Finland. Pregnant women with normal screening results at 12 weeks were approached, and those who provided written informed consent and were proficient in Finnish or Swedish were included. The study was approved by the Ethics Committee of the Hospital District of Southwest Finland (ETMK57/180/2011 and ETMK12/180/2013). Eligible families received study information, and parental consent was obtained in writing. Mothers’ depressive symptoms were assessed at 14, 24, and 34 gestational weeks (GW). Between December 2014 and December 2015, 270 women donated hair samples 1–3 days after childbirth; 230 were eligible for analysis, resulting in 224 samples after outlier exclusion. For brain imaging, infants were excluded if they had perinatal complications, scored less than 5 on the 5-minute Apgar score, had a central nervous system anomaly, underwent a clinical MRI, were born before 32 weeks, or weighed less than 1500 grams at birth. These criteria were confirmed via a structured phone interview, and 180 infants aged 2 to 5 weeks were enrolled. DTI data were obtained from 167 infants, with 37 infants meeting all criteria for the final sample (Figure 2). The study followed Strengthening the Reporting of Observational Studies in Epidemiology (STROBE) guidelines.

**Figure 2.**
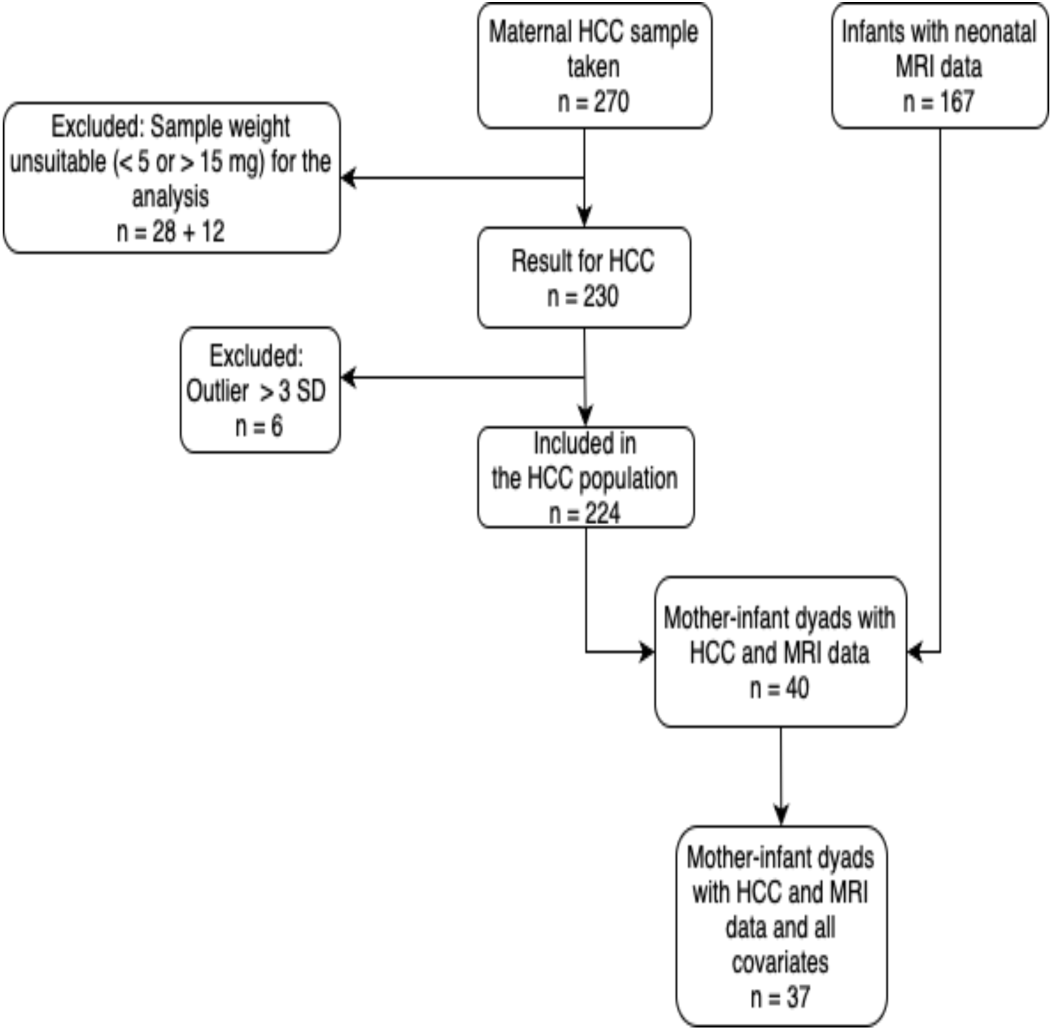
Flowchart of study sample. HCC; maternal hair cortisol concentration. MRI; magnetic resonance imaging.

### 2.2 Infants’ MRI Data Acquisition

For a detailed description of the MRI data acquisition, see Merissari et al. (2019)^37^. Briefly, MRI scans were conducted at the Medical Imaging Centre of the Hospital District of Southwest Finland using a 3T MAGNETOM Skyra scanner (Siemens Healthineers). Infants were fed, swaddled in a vacuum mattress, and no anesthesia was used. They were protected with double hearing protection, while parents, who remained in the room, used standard ear muffs. The session was paused if an infant woke, and a neuroradiologist reviewed the images for incidental findings, reported to families within 1 to 4 weeks. No incidental findings were reported in this sample (see^38^ for overall prevalence). The MRI protocol included diffusion tensor imaging (DTI) with the following parameters: repetition time/echo time = 9300/87.0 ms, field of view = 208 mm, isotropic voxel resolution = 2.0 mm, b value 1000 s/mm², and 96 diffusion directions across three sets. Structural T1- and T2-weighted images and resting-state MRI were also obtained. b0 volumes were visually inspected, and the highest quality volume was selected. Brain masking was performed using FSL’s Brain Extraction Tool^39,40^, and diffusion data quality was assessed with DTIPrep^41^. Poor quality directions were excluded, and the best of the three sequences was used. Motion and eddy current corrections were applied using FSL’s eddy tool^42^, and the data were modeled with FSL’s dtifit.

### 2.3 Image Processing

FA is a composite measure of diffusion directionality and eigenvalues, ranging from 0 (complete isotropic diffusion) to 1 (highly directional diffusion)^43^. We used the Tract-Based Spatial Statistics (TBSS) pipeline from FSL^44^, focusing on WM tract skeletons by aligning individual images to a standardized space. TBSS minimizes alignment issues and partial volume effects by estimating tract centers and does not require spatial smoothing. Preprocessing involved using the “tbss_2_reg -n” option to select the most representative image for creating a study-specific template, which was aligned along the anterior commissure–posterior commissure line. We then used a modified “tbss_3_postreg -S” step to register the data to the template and upsample it to 1 mm³ resolution. Finally, an FA threshold of 0.15 was applied in the “tbss_4_prestats” module to create an FA skeleton. For a detailed description of the preprocessing procedure, see Merissari et al. (2019)^37^.

### 2.4 Prenatal maternal hair cortisol concentration

Maternal hair samples (n=270) were collected 1–3 days post-delivery from a standardized area at the back vertex of the head, closest to the scalp. Each sample consisted of at least 5 mg of hair, with a minimum length of 5 cm, representing cumulative cortisol levels over the preceding 5 months of pregnancy. Hair cortisol extraction was performed at the University of Minho, Portugal, following a protocol adapted from Davenport et al.^45^. Briefly, hair segments were washed with isopropanol, minced, and incubated with methanol at 55°C for 24 hours. After centrifugation, the supernatant was evaporated, and the dried samples were reconstituted with phosphate buffer. Cortisol levels were measured in duplicate using an ELISA kit (IBL International Cortisol Saliva ELISA). Following exclusions (Figure 2), 230 HCC samples were included in the final analyses before outlier examination. For more details, see Mustonen et al., 2019.

### 2.5 Prenatal maternal depressive symptoms

Maternal depressive symptoms during pregnancy were measured using the Edinburgh Postnatal Depressive Scale (EPDS)^46^ at 14, 24, and 34 GW. We combined these measures into a single factor score using a regression-based approach in the lavaan package in R^47^. This method estimates each individual’s score on the latent variable of maternal prenatal depressive symptoms by integrating information from the depressive symptom scores at the three time points.

### 2.6 Covariates

We considered infants’ sex, postmenstrual age (PMA), maternal age, income, and pre-pregnancy body mass index (BMI) as potential confounders. Other relevant descriptives are in Table 1. These data were obtained from birth cohort questionnaires and hospital records kept by The well-being county of Southwest Finland.

**Table 1.** Demographics (n = 37). BMI; Body-mass index, EPDS; Edinburgh Postnatal Depressive Scale, EPDS f1; maternal prenatal depressive symptoms factor score.

| Variable | mean | SD | min | max | SE |
| --- | --- | --- | --- | --- | --- |
| Gestational weeks (GW) | 39.79 | 1.3 | 36.29 | 42 | 0.21 |
| Post menstrual age (PMA) | 43.76 | 1.07 | 42 | 45.71 | 0.18 |
| Infant weight at birth | 3477 | 509 | 2580 | 4670 | 83.68 |
| Infant height at birth | 50.43 | 2.38 | 46 | 56 | 0.39 |
| Apgar 5m | 8.95 | 0.85 | 6 | 10 | 0.14 |
| Maternal BMI | 24.67 | 4.31 | 19.72 | 38.42 | 0.72 |
| Mothers age | 30.11 | 5.05 | 19 | 41 | 0.83 |
| EPDS at 14 GW | 4.75 | 3.58 | 0 | 17 | 0.59 |
| EPDS at 24 GW | 4.35 | 3.34 | 0 | 14 | 0.55 |
| EPDS at 34 GW | 3.7 | 3.18 | 0 | 12 | 0.52 |
| EPDS f1 | 0 | 2.46 | -3.54 | 8.22 | 0.4 |

### 2.7 Data Analysis

Analyses were performed using R^48^ and FSL^40^, including brms^49^ using STAN for implementing Markov chains Monte Carlo (MCMC) sampling^50^, overlapping^51^ and ggplot2^52^ R packages. Statistical analyses were conducted in a Bayesian framework. We first computed bivariate correlations among study variables using Pearson’s r (Figure S1) and computed EPDS factor score. We then adopted a two-steps model comparison approach in order to compare and explore a series of multivariate regression models. Models in both steps were compared in terms of statistical evidence (i.e. support by the data) using information criteria^53^, which enables the evaluation of models considering the trade-off between parsimony and goodness-of-fit^54^. In this regard, it is important to keep in mind that as the complexity of the model increases (i.e. more parameters) the fit to the data increases as well but generalizability (i.e. ability to predict new data) does not necessarily increase^55^. The aim is to find the best balance between fit and generalizability in order to describe, with a statistical model, the important feature of the studied phenomenon, but not the random noise of the observed data. Information criteria provide an estimate of the average deviance (i.e. error) of the model’s ability to predict new data, thus lower values are interpreted as indications of a better model^56^. In the present study, we compared models with the leave-one-out cross-validation information criterion (Loo IC)^57^, where lower values reflect a better fit of the model to data, and the model weight criterion^53^, with higher values reflecting a stronger support for the model. In addition, for each endogenous variable, the variance explained by predictors was explored using the 90% Highest Posterior Density Intervals (HPDI) of R2 on the best model selected^56,58^. HPDI values provide a direct representation of the most credible values of estimated parameters after accounting for prior beliefs.

#### 2.7.1 Prior and Posterior distributions

For prior specifications see Section 2 Supplementary Materials. In summary, given the small sample size, we used weak informative priors: Student’s t(3, 0, 0.01) for infants’ PMA, sex, maternal age and income; Student’s t(3, 0, 0.1) for maternal BMI; Student’s t(3, 0.5, 0.1) for maternal prenatal depressive symptoms; Student’s t(3, -0.1, 0.2) for maternal prenatal HCC; and Student’s t(3, -0.01, 0.01) for the interaction between maternal prenatal depressive symptoms and HCC. Priors incorporated our expectations (mean value) and uncertainty (standard deviation). For example, we expected a weak but negative effect of maternal HCC on brain development, formalized with Student’s t(3, -0.1, 0.2), implying the parameter would fall within [-0.57, 0.37] with 90% probability. We defined a plausible value for the interaction term and the Region of Practical Equivalence^ROPE, 59^ as [-0.005, 0], given the small range of our study variables^60^. If the estimated parameter would fall between this range, the interaction could be considered substantially null. We used MCMC to estimate posterior distributions with four chains of 4000 replicates each, summarizing with median values and the 90% HPDI. We also computed the degree of overlap of the posterior distributions of parameter (eta) ^51^, where eta measures the overlap between two empirical densities, ranging from 0 (no overlap) to 1 (complete overlap). Values within this range quantify the similarity/difference of the values in two groups (e.g., high and low HCC score).

#### 2.7.2 Model comparison

First step, we compared the following models: model 0, the null model (i.e., M00), assuming that there is no relation among study variables, model 1 (i.e., M01), with maternal HCC and EPDS scores predicting infants’ CC FA values, model 2 (i.e., M02), which was similar to model 1 but included the interaction effect between maternal HCC and EPDS scores on infants’ CC FA values to test whether maternal HCC predicted infants’ CC FA values, conditional on maternal EPDS scores. In the second step, once the best model was identified ([M02]), we refined it by adding two crucial variables, infants’ sex and PMA, known for their relevance in the literature. Once these were included, we systematically added additional potentially intervening variables —maternal age ([M02a]), body mass index (BMI) ([M02b]), and income ([M02c])— to [M02] plus infants’ sex and PMA.

#### 2.7.3 Voxel-based statistical analyses

We then perform permutation-based inference on the interaction effects, ensuring robust analysis across the WM regions delineated by the skeleton, using FSL’s randomize tool^61^ and 2D optimized threshold-free cluster enhancement with 5000 permutations^62^. We entered the interaction between maternal prenatal EPDS score and HCC as the explanatory variable in a general linear model and controlled it for the target variables included in best model identified in the model selection (e.g., infant sex and PMA as well as maternal age, BMI, educational level). The models were corrected for multiple comparisons with threshold-free cluster enhancement, at P < .05 across the whole brain FA skeleton.

## 3. Results

### 3.1 Population characteristics

There were 20 female and 17 male infants. Maternal education levels varied: 12 infants’ mothers had primary education, 15 had secondary education, and 10 had tertiary education. Most infants were from families with moderate incomes: 14 from the lowest income bracket, 19 from the middle bracket, and 4 from the highest bracket. ICU admission was rare, with only 1 infant requiring ICU care. Smoking during the 1st trimester was reported in 3 cases, but no mothers reported alcohol or drug use. No maternal complications were reported (Section S1).

### 3.2 Multivariate regression models

Table 2 and 3 show the results of Step 1 and 2 of the Bayesian model comparison. Regarding Step 1, the best performing model was model M02, with maternal HCC interacting with EPDS factor score in predicting infants’ Genu, Body and Splenium FA values. Regarding the second model comparison, the best performing model was the one with infants’ sex, PMA and maternal age as covariates (M02a, a detailed description of model comparison is provided in Section S3). We describe the best model in detail below and its coefficients are reported in the Supplementary Material, Box 2.

**Table 2.** Model Comparison Step 1. M00 is the null model, M01 is the model with the additive role of EPDS and maternal HCC predicting CC FA values, M02 includes the interaction term between EPDS and maternal HCC on CC FA values. The best fitting model for is the first one. Models are estimated on N=37 subjects. LOO; leave-one-out cross-validation information criterion, W; model weight.

| Model | LOO | SE | W |
| --- | --- | --- | --- |
| M02 | -527.3 | 15.8 | 0.593 |
| M01 | -526.3 | 16.1 | 0.36 |
| M00 | -522.1 | 17.54 | 0.045 |

**Table 3.**
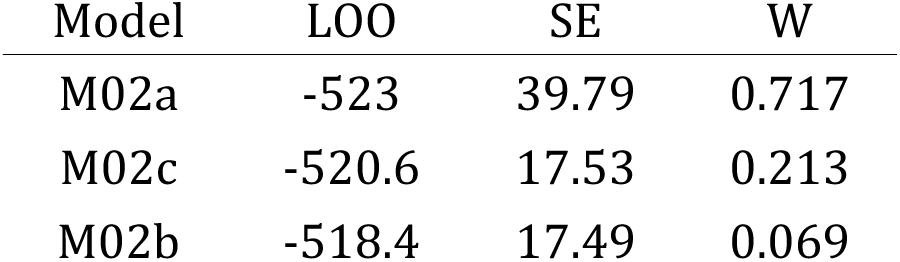
Model Comparison Step 2. All the models here are controlled for infants’ sex and post-menstrual age. M02a is the model with maternal age, M02b is the model with maternal BMI, M02c includes maternal income. The best fitting model is the first one. Models are estimated on N=37 subjects. LOO; leave-one-out cross-validation information criterion, W; model weight.

| Model | LOO | SE | W |
| --- | --- | --- | --- |
| M02a | -523 | 39.79 | 0.717 |
| M02c | -520.6 | 17.53 | 0.213 |
| M02b | -518.4 | 17.49 | 0.069 |

We analyzed model predictions based on the parameters’ posterior distributions and the Posterior Predictive Distribution (PPD), i.e., the distribution of possible unobserved values conditional on observed data and model parameters^63^. Credible intervals of the explained variance for each exogenous variable did not include the zero for the best model selected, with R2 90% HPDI (0.21, 0.55) for Genu, R2 90% HPDI (0.17, 0.51) for Body and R2 90% HPDI (0.15, 0.50) for Splenium, suggesting that the effects can be reasonably supported. More specifically, posterior distributions and associated 90% HPDI showed that the interaction between EPDS scores and maternal HCC has a negative association on infants’ FA considering all the three sections of the infants’ CC [B_Genu_= -0.003, 90% HPDI (-0.005, -0.001); B_Body_ = -0.003, 90% HPDI (-0.006, -0.001); B_Splenium_ = -0.004, 90% HPDI (-0.008, -0.001)]. No association of infants’ sex with CC FA values were identified [B_Genu_= -0.001, 90% HPDI (-0.008, 0.008); B_Body_ = 0.005, 90% HPDI (-0.006, 0.014); B_Splenium_ = 0.003, 90% HPDI (-0.010, 0.017)]. Similarly, no association emerged between infants’ CC FA values and maternal age [B_Genu_= -0.001, 90% HPDI (-0.001, 0.006); B_Body_ = 0.001, 90% HPDI (-0.001, 0.002); B_Splenium_ = 0.002, 90% HPDI (-0.001, 0.003)]. Differently, infants’ PMA resulted in positive association with CC FA values of Genu and Splenium [B_Genu_= 0.005, 90% HPDI (0.001, 0.010); B_Splenium_ = 0.009, 90% HPDI (0.001, 0.016)] but not with CC Body [B_Body_ = 0.003, 90% HPDI (-0.003, 0.007)]. For further details, Section 3 of the Supplementary Materials provides diagnostic plots (Figure S6), Posterior Predictive Check (Figure S7) and the comparison between prior and posterior distributions of the interaction parameter, i.e., ROPE (Figure S9). Finally, Figure 3 illustrates the overlapping index of M01 PPD or, in other words, of the expected values of infants’ CC FA values as a function of EPDS scores (x-axis) and maternal HCC (colors). The values presented in the plot quantify the variation in infants’ FA values based on maternal depressive symptoms in groups stratified by high (1 SD above the mean) and low (1 SD below the mean) HCC levels, where maternal HCC is expressed logarithmically and maternal depressive symptoms are a factor score. For example, regarding infants’ CC Splenium FA values, infants from mothers with low (1.73) versus high (3.93) maternal HCC levels showed a substantial overlap (0.76) when EPDS scores were low (-3.54), indicating minimal differences in means. Conversely, those with higher EPDS scores (0.53) exhibited a smaller overlap (0.49). When EPDS scores reached the maximum (8.22), the overlap index between infants from mothers with high versus low HCC levels was very low (0.11), suggesting relevant differences in means. Results from the whole-brain regression model confirmed our results (Figure S10).

**Figure 3.**
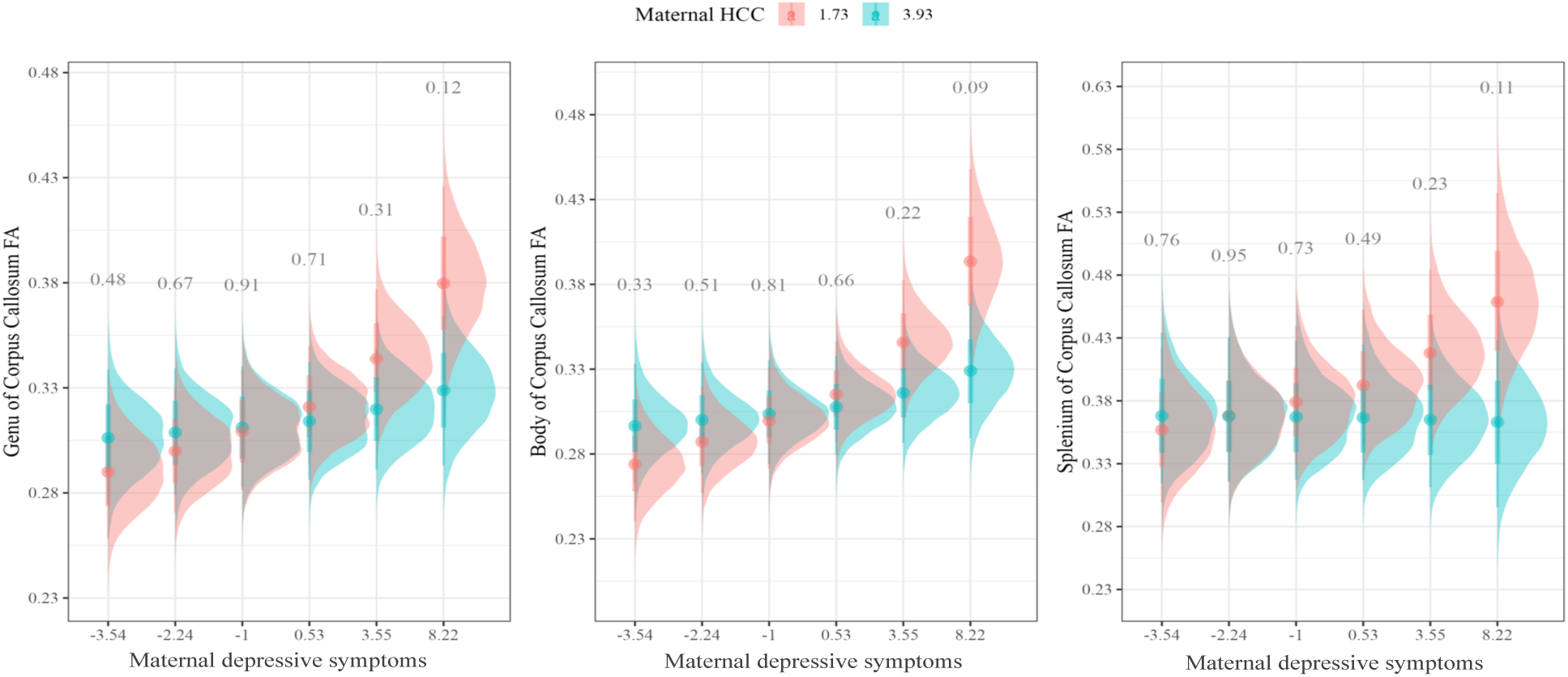
Densities represent conditional posterior distributions for mothers with high HCC (1 SD above the mean, azure densities) and low HCC (1 SD below the mean, pink densities). The values reported are the overlapping indices (eta) between pairs of densities. Maternal HCC is expressed in logarithms; maternal depressive symptoms are a factor score.

## 4. Discussion

The study explored the relationship between maternal prenatal depressive symptoms, HCC, and diffusion properties (i.e., FA) of the CC in newborn brain MRIs. We found a negative association between maternal depressive symptoms and HCC in predicting infants’ CC FA. Specifically, higher prenatal depressive symptoms were linked to higher infants’ FA when maternal HCC values were low (1 SD below the mean). Additionally, a positive association was observed between infants’ PMA and FA values of the Genu and Splenium of the CC. Most associations were modest, consistent with the complex effects of prenatal environment on the brain. However, some associations, particularly in the Splenium, were stronger, with the posterior 30% outside the ROPE. These findings are novel, as there are no previous reports linking prenatal maternal depressive symptoms, HCC, and neonates’ WM integrity.

The interaction between HCC and prenatal depressive symptoms was negatively associated with FA in the CC Genu, Body, and Splenium. Higher prenatal depressive symptoms were linked to higher FA only when HCC values were 1 SD below the mean, suggesting that increased maternal depressive symptoms strengthen the association with neonates’ CC integrity at lower maternal HCC levels, with no association for high HCC levels. These findings are somewhat unexpected but align with previous literature suggesting a potential protective role of maternal cortisol against the adverse effects of psychopathology on infant brain development^64–66^. Cortisol levels naturally increase towards the end of pregnancy; it rises up to 2–3-fold to prepare both the mother and the fetus for parturition^67^. This increase ensures that the fetus is exposed to sufficient cortisol during the third trimester, which is crucial for the maturation of the fetal lungs and the preparation for delivery^68,69^. A less pronounced increase could represent a suboptimal environment, explaining observations where higher end-of-pregnancy cortisol levels were associated with improved motor, cognitive, behavioral and emotional development^64–66^. The diminished exposure to maternal prenatal cortisol levels, especially simultaneously with other signals of increased maternal depressive symptoms, could lead to boosted offspring WM maturation. These results are also in line with the stress acceleration hypothesis^70^; the combined effect of maternal prenatal depressive symptoms and atypical HCC concentration serve as early stressors that promotes accelerated fetal brain development.

Our results also support the idea that experienced psychological distress and altered HPA axis activity are interrelated but distinct phenomena, with maternal depressive symptoms affecting fetal programming through multiple pathways, only some of which are directly linked to maternal prenatal cortisol levels^29,71^. Maternal depressive symptoms have been associated with the downregulation of placental 11β-HSD2 activity in metabolizing cortisol into inactive cortisone^3^, and with epigenetic alterations, such as elevated methylation of 11β-HSD2 and NRC31, a glucocorticoid receptor gene^72,73^. These findings highlight the complexity of cortisol regulatory systems during pregnancy and early development, emphasizing the need to assess HPA axis functioning alongside other measures of psychosocial stress.

Furthermore, our study unveiled a positive association between PMA and FA values in the CC’s Genu and Splenium, but not in the Body. This suggests that the CC is non-uniformly affected by PMA. During early life, the brain undergoes significant changes in WM microstructure, including increased volume, myelination and axonal density. FA values, reflecting the organization and coherence of WM tracts, typically rise with age, indicating tract maturation. However, FA growth is not linear and varies across brain regions. Our results align with previous studies showing that different CC regions exhibit distinct development patterns^74^.

We did not test the behavioral phenotype in association with our results, which could represent a relevant step forward. Mustonen et al. (2024) found that, in two-year-old children, the interaction between maternal late pregnancy HCC and prenatal depressive symptoms was negatively associated with internalizing and dysregulation problems. Borchers et al. (2021) found that higher levels of prenatal depressive symptoms were linked to higher FA of the CC Genu, which was positively associated with behavioral problems at 18 months. Another study found that maternal postnatal depressive symptoms were positively correlated with infant negative reactivity in infants with high FA in the CC and cingulum^22^. Notably, Nolvi (2020), Mustonen (2024), and the present study are derived from the same cohort but different samples, consistently pointing to the effects of maternal prenatal distress on infant outcomes, which strengthens our findings. This evidence suggests that the CC is sensitive to prenatal maternal depressive symptom exposure and HCC atypical concentrations and could represent a neural endophenotype linking early-life stress and offspring less-than-optimal behavioral phenotype. However, this interpretation remains speculative and is an objective for future studies.

As we aimed to explore the cumulative effect of less-than-optimal maternal well-being during pregnancy on offspring brain development, we collapsed three different time measurements of maternal depressive symptoms into one variable. Future studies should consider assessing maternal depressive symptoms at specific gestational periods.

Methodologically, our study benefited from robust techniques for CC delineation in DTI data and the utilization of the tensor model, which proved suitable for analyzing CC data^35^. Concerning HCC, we used 5 cm segments instead of the typical 3 cm segments, which may have diluted some pregnancy-specific alterations in HCC^66,75^. This choice was partly based on practical reasons to ensure adequate sample weight for reliable analyses. Overall, the risk of Type II (false negative) rather than Type I (false positive) error may have increased.

The present study has some limitations, including a small sample size with DTI and HCC data and participant population homogeneity, which may impact the generalizability of our findings. Future studies should involve larger, more diverse cohorts. Additionally, this study did not include genetic measures, which could play a relevant role in the interplay between prenatal maternal well-being, HCC, and offspring WM maturation. Genetic and epigenetic factors, including cortisol’s potential as an epigenetic modifier, should be considered in future research. Finally, while the narrow range of EPDS scores in this non-clinical population calls for cautious interpretation, it should be noted that the association between HCC and CC FA appears evident even at these moderate symptom levels.

In conclusion, our study provides valuable insights into the complex interplay between maternal prenatal cortisol levels, depressive symptoms, and infant CC integrity. By unraveling these dynamics, we aim to contribute to a deeper understanding of the mechanisms linking maternal depressive symptoms and late-pregnancy cortisol concentrations to offspring brain structure. Importantly, our findings highlight the need to consider not only lower FA values but also the potential risks associated with elevated FA, which has already been shown to be problematic in pediatric populations^24,76,77^. This emphasizes the multifaceted nature of neurodevelopmental outcomes and the importance of a nuanced approach in assessing early brain development. Further investigations are required to identify the prognostic relevance of early-life CC development in offspring born to mothers with high levels of depressive symptoms and low cortisol concentration during the end of pregnancy.

## Acknowledgments and Disclosures

JJT, HK, LK, SK, and PM conceptualized and planned the study. SK and PM participated in the data collection. SK, PM, and BC performed the hair sample preprocessing. BC and AJR performed the HCC analyses. JJT designed the data processing pipelines and supervised ILCMW. The original draft was written by ILCMW, reviewed and edited by PM, LK, SN, SK, NMS, MP, KT, BC, AJR, NS, HK and JJT. Statistical analysis and visualizations were done by ILCMW and MP. Project administration was done by LK and HK. Funding was provided by JJT, HK and LK. All authors commented and accepted the final version of the manuscript. ILCMW was supported by a postdoctoral fellowship from the Sigrid Juselius Foundation. PM was supported by the Päivikki and Sakari Sohlberg’s Foundation. SK was supported by Yrjö Jahnsson foundation #20227531. SN and KT were supported by The Research Council of Finland, Centre of Excellence in Learning Dynamics and Intervention Research (#346121). LK was supported by the Research Council of Finland #308176 and #342748, Signe and Ane Gyllenberg Foundation, Finnish State Grants for Clinical Research, Brain and Behavior Research Foundation, Young Investigator Grant #1956, Yrjö Jahnsson Foundation #6976 and #6847, Strategic Research Council (SRC) established within the Academy of Finland #352648; subproject 352655. JJT was supported by the Finnish Medical Foundation, the Emil Aaltonen Foundation, the Sigrid Juselius Foundation, the Signe and Ane Gyllenberg Foundation, the Hospital District of Southwest Finland State Research Grants, the Alfred Kordelin Foundation, the Juho Vainio Foundation and the Orion research Foundation. The authors report no biomedical financial interests or potential conflicts of interest.

