## Supplementary Material for "Association between maternal depressive symptoms and hair cortisol concentration during pregnancy with corpus callosum integrity in newborns"

September 6, 2024

**Abstract**

Leveraging data from the FinnBrain Birth Cohort Study, we analyzed data from 37 mother-infants dyads. MRI data were obtained on 2-5-week-old infants and DTI imaging was conducted to estimate fractional anisotropy (FA) values in CC regions (Genu, Body, and Splenium). We evaluated maternal prenatal cortisol levels through hair cortisol concentration (HCC) extracted from a 5cm hair segment. The latter reflects cortisol concentration over the last five months of pregnancy. To create a comprehensive measure of maternal depressive symptoms during pregnancy, a factor score was computed from EPDS questionnaire data gathered at gestational weeks 14, 24, and 34. We used multivariate regression models adopting a Bayesian approach for statistical testing and controlled our models for several maternal and infant related attributes. Our results revealed that maternal prenatal depressive symptoms and HCC negatively interact in predicting infants' FA in all the CC regions considered. Infants exposed to high prenatal maternal depressive symptoms and low HCC (i.e., HCC score 1 SD below the mean), showed higher FA in the CC [B Genu = -0.003, 90% HPDI (-0.005, -0.001); B Body = -0.003, 90% HPDI (-0.006, -0.001); B Splenium = -0.004, 90% HPDI (-0.008, -0.001)].

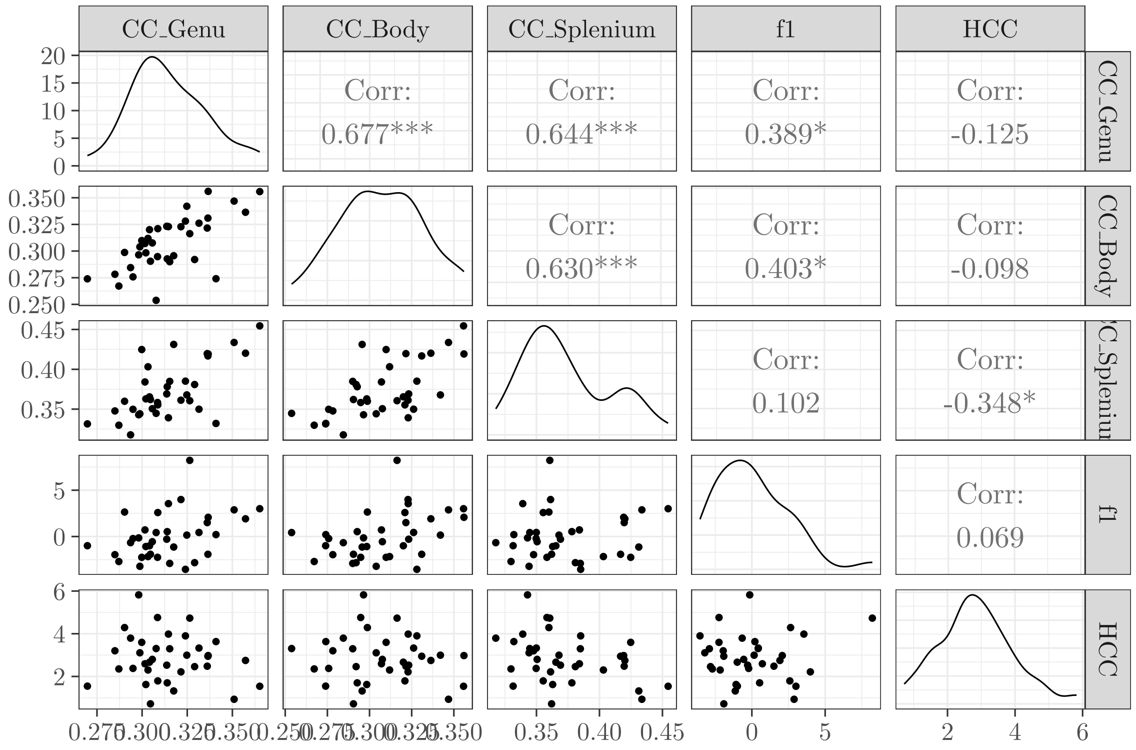

Figure S1: Univariate densities and bivariate distributions among the target variables. CC; corpus callosum, f1; maternal prenatal depressive symptoms (e.g., EPDS factor score), HCC; maternal prenatal hair cortisol concentration.

### 1 Descriptives

The data for maternal monthly income estimate, alcohol use, and illicit drug use are from questionnaires at gestational week 14. The pregnancy complications include a diagnosis (according to ICD-10) for O12 (Gestational edema and proteinuria without hypertension), O13 (Gestational hypertension without significant proteinuria), O14 (Severe pre-eclampsia), O24 (Diabetes mellitus in pregnancy, childbirth, and the puerperium), O46 (Antepartum hemorrhage, not elsewhere classified), or O99.0 (Anemia complicating pregnancy, childbirth and the puerperium). In Figure S1, the empirical distributions of key variables are represented. Note: HCC variables are expressed in logarithms, and the variable `f1` is a factor score.

### 2 Prior Specification

#### 2.1 Prior estimation for interaction term

To estimate interaction prior we used the following model:

$$\text{CC\_genu} = \beta_0 + \beta_1 \text{f1} + \beta_2 \text{HCC} + \beta_3(\text{f1} \cdot \text{HCC}) + \epsilon \tag{1}$$

The first step was to establish plausible values for $\beta$ to define meaningful priors and a *Region of Practical Equivalence*. For simplicity, we centered the dependent variable:

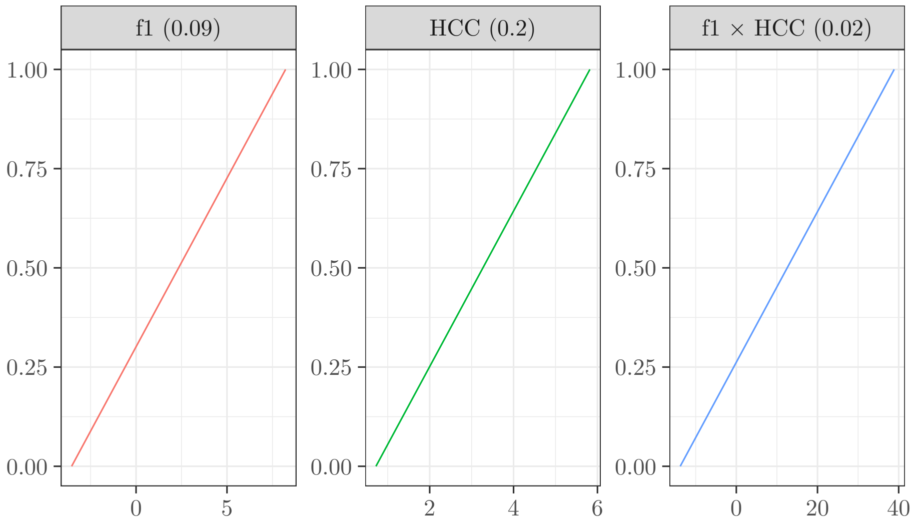

Figure S2: Geometric representation of lines with maximum positive slope with values of $y$ in the range $[0, 1]$ and values of $x$ within the empirical intervals of the variables f1 [-3.54, 8.22], HCC [0.72, 5.82] and f1 imes HCC. The values in parentheses are the regression coefficients of the three lines. The process holds even when considering negative coefficients.

```
> X$CC_Genu.c <- X$CC_Genu - mean( X$CC_Genu )
```

In this way, the intercept becomes zero, allowing us to focus on three parameters instead of four.

First of all, we defined the Design Matrix (DM), $\mathbf{X}$, with a set of representative values for the predictors. For `f1`, we chose $(-3, 3)$, and for `HCC`, $(0, 0.67, 1.33, 2, 2.67, 3.33, 4, 4.67, 5.33, 6)$. The values of `f1` - being factor scores, i.e., standardized scores - represent values for subjects with depressive symptoms $\pm 3$ standard deviations from the mean. This choice is based on the expectation that the majority of subjects in the population fall within this range. The values of `HCC` have been chose based on the empirical range, and for simplicity, we transformed them into deviations from the mean.

To complete the DM, we added a column of 1 (representing the intercept) and a column given by the product `f1` $\times$ `HCC`, representing the interaction. In practice, the DM consists of 20 rows (the combination of selected values) and four columns. As an example, we reported the first few rows of DW below.

```
  Intercept f1       HCC f1.HCC
1         1 -3 -3.000000      9
2         1  3 -3.000000     -9
3         1 -3 -2.333333      7
4         1  3 -2.333333     -7
```

#### 2.2 Looking for $\beta$ plausible values

First, we considered a purely geometric rationale. In a linear model, the coefficient $\beta$ expresses the difference in values on $Y$ for each unit difference in $X$. Since, in this case, the dependent variable

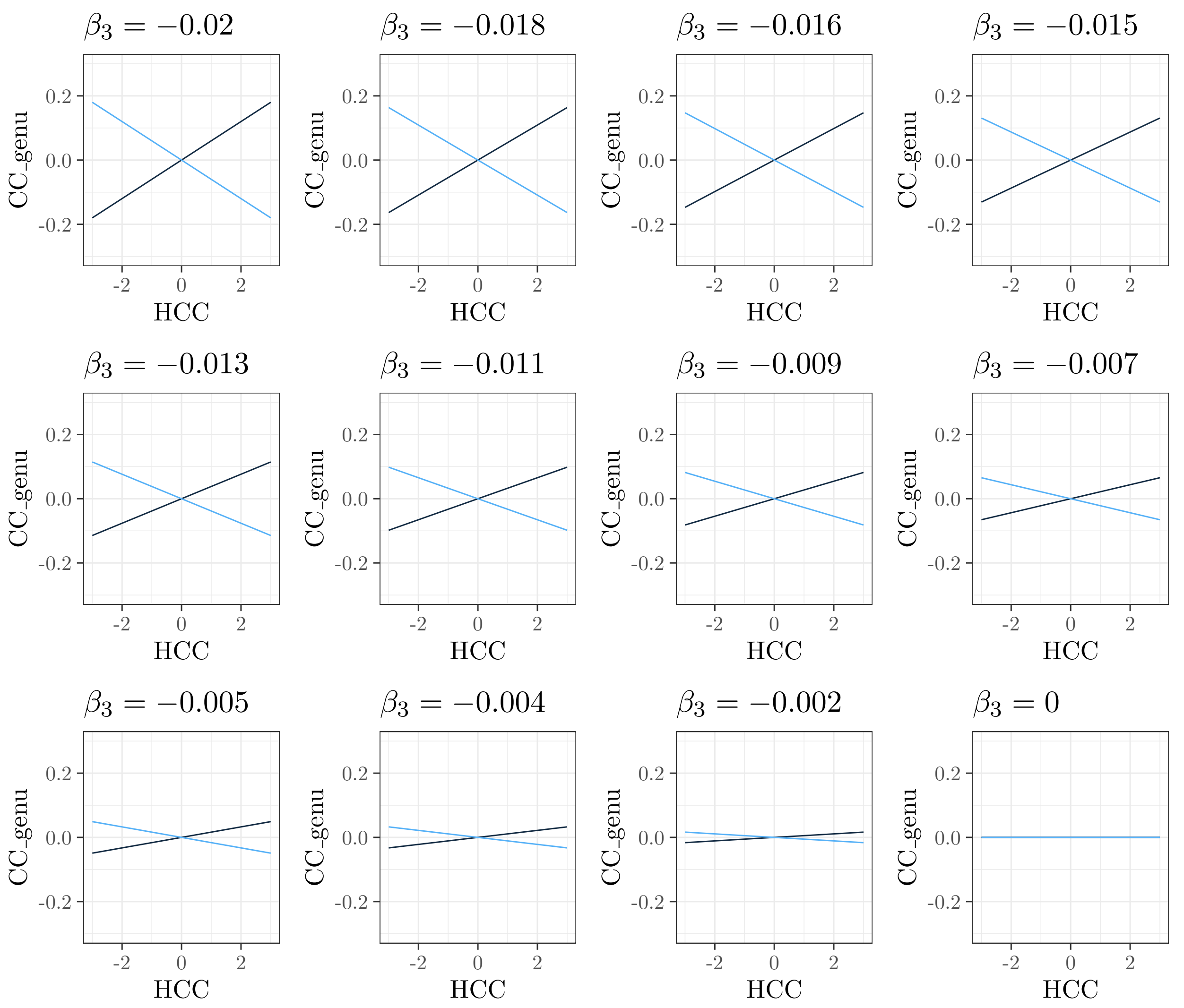

Figure S3: Examples of interaction between HCC (on the x-axis) and f1; the light color is associated with the condition f1 = -3, and the dark color is associated with the condition f1 = 3.

CC_genu theoretically takes values in the interval $[0, 1]$—in our sample $[0.27, 0.37]$— it means that the parameters $\beta_1$ and $\beta_2$ will presumably be less than 1. In particular, as the empirical range of $1 \in [-3.54, 8.22]$, the parameter $\beta_1$ will plausibly be less than $1/12 = 0.09$. Similarly, since the empirical range of HCC $\in [0.72, 5.82]$, the parameter $\beta_2$ will probably be less than $1/5 = 0.2$.

In Figure S2, the left and center panels represent lines with the maximum slope concerning `f1` and `HCC`, respectively. Each of these lines holds when the other $\beta$ is fixed at zero. In the third panel, on the right, the line associated with the maximum possible value of interaction is represented, again with the constraint that the other $\beta$ values are zero. It should be clear that the same consideration holds if the parameters are negative, in the sense that they would be equal in absolute value.

##### 2.2.1 Exploring the interaction between `f1` and `HCC`

To assess the interaction graphically, we considered the simplest case: all variables centered, and parameters $\beta_1$ and $\beta_2$ fixed at zero. Therefore, we choose 12 values for the interaction parameter

$\beta_3 \in [-0.02, 0]$ and calculate the expected values.

For example, if we take the first value, $\beta_3 = -0.02$, we can obtain a vector of expected values for CC_genu based on the equation:

$$\hat{Y} = 0 \times \text{f1} + 0 \times \text{HCC} + -0.02 \times (\text{f1} \times \text{HCC}) \tag{2}$$

In the top-left panel of Figure S3, the expected values obtained are represented; in the other panels, all interactions obtained based on the chosen values of $\beta_3$. On the x-axis, there are the (centered) values of HCC, and on the y-axis, there are the expected values, which, being centered, will always fall between -0.5 and 0.5. The two colors are related to the two values of f1, light for -3 and dark for 3. Based on this figure, we can identify a value of $\beta_3$ below which the interaction can be considered *not relevant*, and consequently, define the Region of Practical Equivalence (ROPE). We established that the ROPE is $[-0.005, 0]$, in other words, interaction values falling within this interval were considered substantially null.

##### 2.3 Prior distribution

For each models parameter, we defined a prior probability distribution, chosen with the aim of formalizing our prior hypotheses. In particular, considering that the sample size is small, we used the following weak informative priors: for infants' post-menstrual age (PMA) and sex as well as maternal age and income (on CC genu, body and splenium) effect Student's t(3, 0, 0.01), for maternal BMI (on CC genu, body and splenium) effect Student's t(3,0, .1), for maternal prenatal depressive symptoms effect (on CC genu, body and splenium) Student's t(3, 0.5, 0.1), for maternal prenatal hair cortisol concentration effect (on CC genu, body and splenium) Student's t(3, -0.1, 0.2), and for interaction effect between maternal prenatal depressive symptoms and hair cortisol concentration Student's t(3, -0.01, 0.01). In this regard, we defined prior distributions to incorporate our expectations (defined by prior mean value) and associated uncertainty (defined by prior standard deviation) into the analysis. Figure S4 represents prior distributions. Box 1 reports in detail all the priors. The "coef" column stands for the model coefficient names, the "prior" column for priors' degrees of freedom, mean and sd, the "class column for parameter for which the prior is specified (b is the coefficients (slopes) of the predictors in the model) and the "response" is for the response variable. Credible Interval column shows corresponding intervals within which we hypothesized that model parameters would fall with a 90% of probability.

#### 3 Model comparison

A detailed description of the strategy adopted in the present work is already provided in the main text file. However, highlighting only the essential points, we used a model-comparison strategy to identify the best model based on the following performance indices: the Leave-One-Out cross-validation Information Criterion (LOO; Vehtari, Gelman, & Gabry, 2017) the Bayesian R2 (Gelman, Goodrich, Gabry, & Vehtari, 2019), and the model weights (w; Yao, Vehtari, Simpson, & Gelman, 2018). Lower values of LOO and higher values of w indicate a more plausible model.

First step: we compared the following models: model 0, the null model (i.e., M00), i.e. a model assuming that there is no correlation among study variables, model 1 (i.e., M01), with maternal HCC and EPDS scores predicting infants' CC FA values, model 2 (i.e., M02), which was similar to model 1 but included the interaction effect between maternal HCC and EPDS scores on infants' CC FA values to test whether maternal HCC predicted infants' CC FA values, conditional on maternal EPDS scores.

```
                       prior class            coef       resp
1      student_t(3, 0.5, 0.1)     b              f1     CCBody
2     student_t(3, -0.1, 0.2)     b             HCC     CCBody
3   student_t(3, -0.01, 0.01)     b          f1:HCC     CCBody
4         student_t(3,0,0.01)     b        SEXtytto     CCBody
5         student_t(3,0,0.01)     b KOKONAISIKA_VKO     CCBody
6          student_t(3,0,0.1)     b    maternal_BMI     CCBody
7            student_t(3,0,1)     b     mothers_age     CCBody
8            student_t(3,0,1)     b maternal_income     CCBody
9        student_t(3, 0, 0.1) sigma                     CCBody
10     student_t(3, 0.5, 0.1)     b              f1     CCGenu
11    student_t(3, -0.1, 0.2)     b             HCC     CCGenu
12  student_t(3, -0.01, 0.01)     b          f1:HCC     CCGenu
13       student_t(3, 0, 0.1) sigma                     CCGenu
14        student_t(3,0,0.01)     b        SEXtytto     CCGenu
15        student_t(3,0,0.01)     b KOKONAISIKA_VKO     CCGenu
16         student_t(3,0,0.1)     b    maternal_BMI     CCGenu
17           student_t(3,0,1)     b     mothers_age     CCGenu
18           student_t(3,0,1)     b maternal_income     CCGenu
19     student_t(3, 0.5, 0.1)     b              f1 CCSplenium
20    student_t(3, -0.1, 0.2)     b             HCC CCSplenium
21  student_t(3, -0.01, 0.01)     b          f1:HCC CCSplenium
22        student_t(3,0,0.01)     b        SEXtytto CCSplenium
23        student_t(3,0,0.01)     b KOKONAISIKA_VKO CCSplenium
24         student_t(3,0,0.1)     b    maternal_BMI CCSplenium
25           student_t(3,0,1)     b     mothers_age CCSplenium
26           student_t(3,0,1)     b maternal_income CCSplenium
27       student_t(3, 0, 0.1) sigma                 CCSplenium
```

Box 1: Priors. coef = model coefficient names; prior = priors' degrees of freedom, mean and sd; class = parameter for which the prior is specified (b is the coefficients (slopes) of the predictors in the model); resp = response variable; f1 = maternal prenatal depressive symptoms (e.g., EPDS factor score), HCC = maternal prenatal hair cortisol concentration; f1 $\times$ HCC = interaction between maternal prenatal depressive symptoms and maternal prenatal hair cortisol concentration; SEXtytto = female infants; KOKONAISIKA_VKO = infants post-menstrual age.

Second step: after identifying the best model ([M02]), we refined it by adding two crucial variables, infants' sex and PMA, known for their relevance in the literature. Once these were included, we systematically added additional potentially intervening variables —maternal age ([M02a]), body mass index (BMI) ([M02b]), and income ([M02c])— to this enhanced model (i.e., [M02] plus infants' sex and PMA). A visual representation of this two-step model comparison process is provided in Figure S5.

### 4 Results

For the model M02a, which resulted as the best model (i.e. the model with the highest w), we reported below:

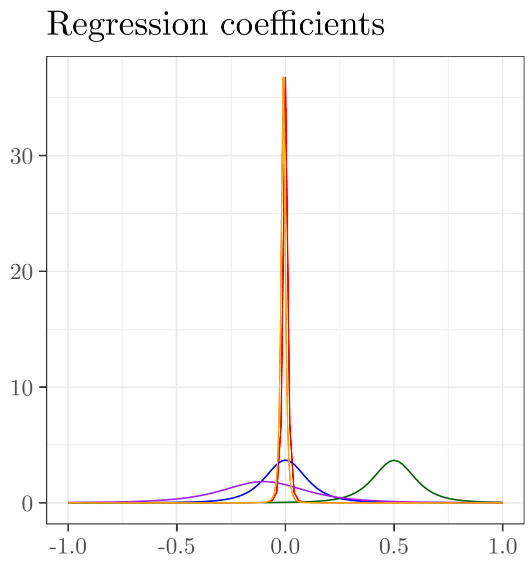

Figure S4: Prior distribution. Red = infants' post-menstrual age and sex and maternal age and income; blue = maternal BMI; dark green = maternal prenatal depressive symptoms; purple = maternal prenatal hair cortisol concentration; orange = the interaction term between maternal prenatal depressive symptoms and hair cortisol concentration. All these effects are on CC genu, body and splenium.

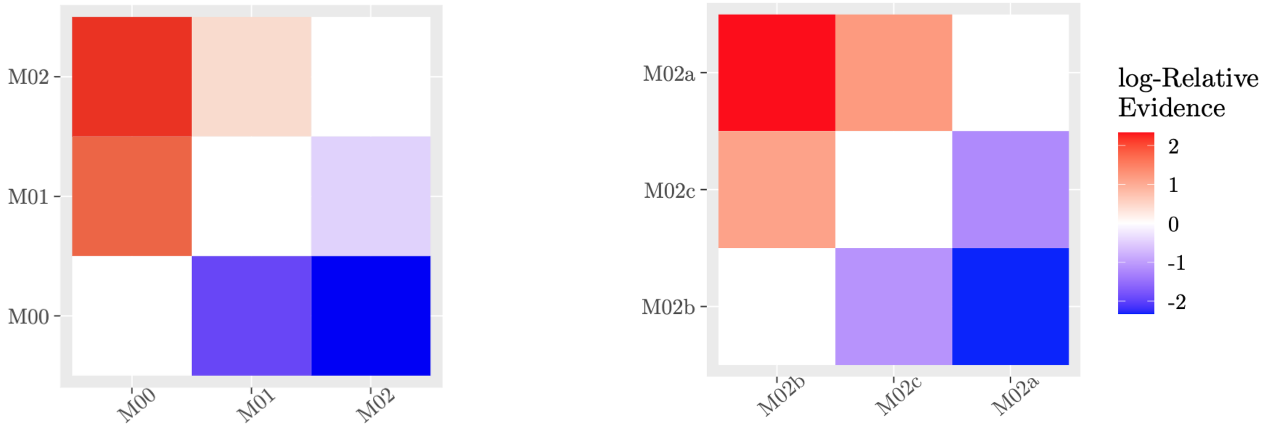

Figure S5: Two-steps model comparison; the left panel refers to the first step while the right panel refers to the second step.

(i) three diagnostics plots representing, respectively: residuals vs fitted values (Residuals vs Fitted plot); normality of residuals (Normal QQ-plot); and square root of absolute values of standardized residuals vs fitted values (Scale-location plot).

(ii) posterior predictive check plot, i.e., graphical comparison between data simulated from the posterior predictive distribution and real-world observations.

(iii) model coefficients.

(iv) Comparison between prior and posterior distributions of interaction parameters.

#### 4.1 Diagnostics plots

Figure S6 reports three diagnostics plots representing residuals vs fitted values (Residuals vs Fitted plot); normality of residuals (Normal QQ-plot); and square root of absolute values of standardized residuals vs fitted values (Scale-location plot). The hallmarks of a proficient model include residuals that are independent from the fitted values (as depicted in the Residuals vs Fitted plot), a normal distribution (as illustrated in the Normal QQ-plot), and a consistent variance in relation to the fitted values (as shown in the Scale-location plot).

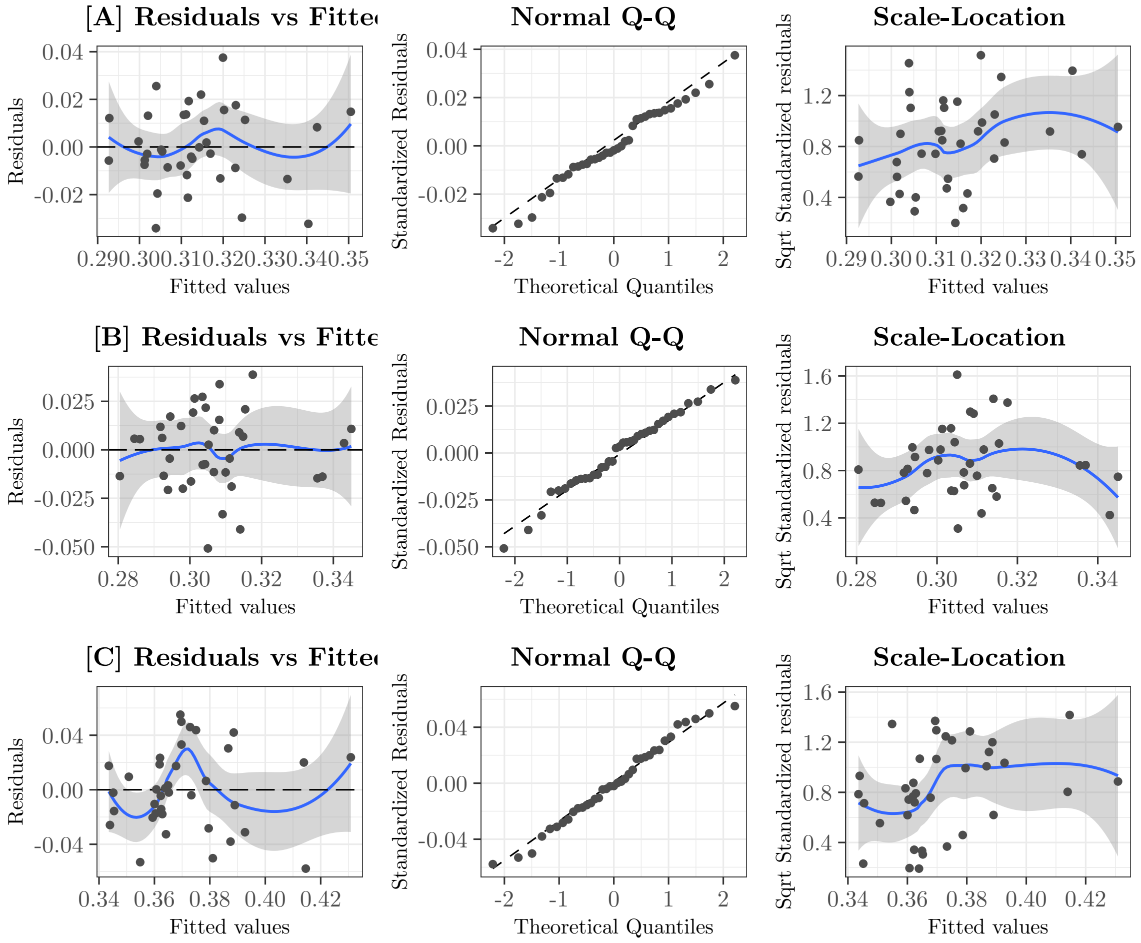

Figure S6: Diagnostic plots of the target model [M02a] .

#### 4.2 Posterior Predictive Check

The posterior predictive check (PPC) generates replicated data following the posterior predictive distribution. Figure S7 report PPC of the target model considering CC Genu [A], Body [B] and Splenium [C] as outcome variables. If a model has a good fit, the generated data (ligh blue lines) looks like the observed data (dark blue line) (Gabry, Simpson, Vehtari, Betancourt, & Gelman, 2019).

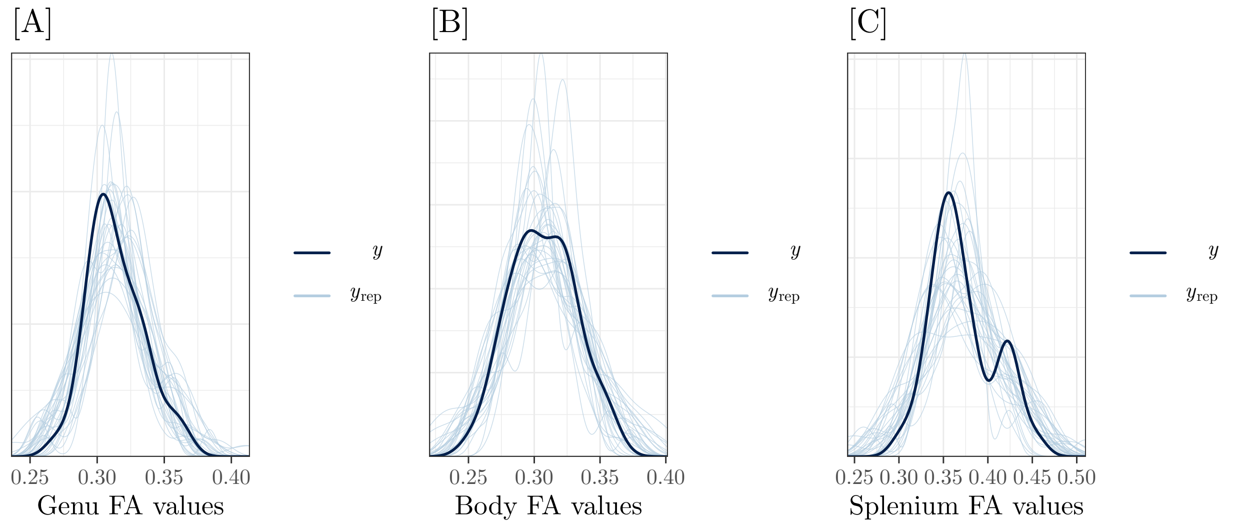

Figure S7: Posterior Predictive Check of the best model [M02a] considering Corpus Callosum Genu [A], Body [B] and Splenium [C] as outcome variables.

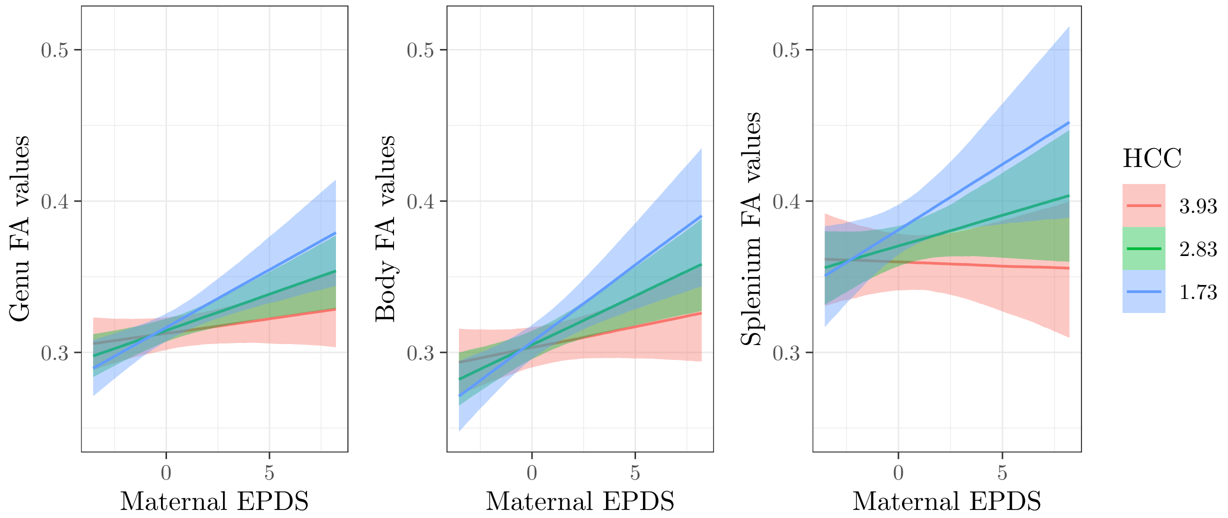

Figure S8: Interaction plots. Expected values of Corpus Callosum Genu, Body and Splenium as a function of the interaction HCC and f1; the values selected for f1 are the mean ± the standard deviation.

##### 4.3 Model coefficients

Box 2 report models coefficients while Figure S8 summarizes the predictions on `CC_Genu`, `CC_Body` and `CC_Splenium` based on the interaction `HCC` × `f1` are displayed.

##### 4.4 Prior vs Posterior

In Figure S9, prior is represented by the blue line, while the posterior is the red area. Considering CC Genu, the posterior is almost entirely within the ROPE — specifically, 98% of the distribution lies in the interval [-0.005, 0] — supporting the idea that the interaction, while evident, is not very

```
 Family: MV(gaussian, gaussian, gaussian)
  Links: mu = identity; sigma = identity
         mu = identity; sigma = identity
         mu = identity; sigma = identity
Formula: CC_Genu ~ f1 * HCC + SEX + KOKONAISIKA_VKO + mothers_age
         CC_Body ~ f1 * HCC + SEX + KOKONAISIKA_VKO + mothers_age
         CC_Splenium ~ f1 * HCC + SEX + KOKONAISIKA_VKO + mothers_age
   Data: data (Number of observations: 37)
  Draws: 4 chains, each with iter = 2000; warmup = 1000; thin = 1;
         total post-warmup draws = 4000

Population-Level Effects:
                           Estimate Est.Error l-90% CI u-90% CI  Rhat Bulk_ESS Tail_ESS
CCGenu_Intercept              0.101     0.119   -0.090    0.300 1.000     3904     2889
CCBody_Intercept              0.179     0.145   -0.053    0.418 1.001     4038     2958
CCSplenium_Intercept         -0.056     0.206   -0.397    0.278 1.001     3420     3161
CCGenu_f1                     0.012     0.004    0.006    0.018 1.002     2813     2985
CCGenu_HCC                   -0.002     0.003   -0.007    0.003 1.001     3126     2808
CCGenu_SEXtytto              -0.000     0.005   -0.009    0.009 1.001     3746     2797
CCGenu_KOKONAISIKA_VKO        0.005     0.003    0.001    0.010 1.001     3743     3016
CCGenu_mothers_age           -0.000     0.001   -0.002    0.001 1.000     4843     3461
CCGenu_f1:HCC                -0.003     0.001   -0.004   -0.001 1.002     2914     3268
CCBody_f1                     0.016     0.005    0.009    0.023 1.001     2928     2593
CCBody_HCC                   -0.002     0.004   -0.008    0.004 1.000     3450     3130
CCBody_SEXtytto               0.005     0.006   -0.006    0.015 1.001     3557     2559
CCBody_KOKONAISIKA_VKO        0.003     0.003   -0.003    0.008 1.001     3902     3326
CCBody_mothers_age            0.000     0.001   -0.001    0.002 1.000     5404     3578
CCBody_f1:HCC                -0.003     0.001   -0.005   -0.001 1.001     2994     2767
CCSplenium_f1                 0.017     0.007    0.006    0.028 1.002     2604     2890
CCSplenium_HCC               -0.010     0.005   -0.018   -0.001 1.000     3124     3114
CCSplenium_SEXtytto           0.003     0.008   -0.010    0.017 1.000     3111     2516
CCSplenium_KOKONAISIKA_VKO    0.009     0.005    0.002    0.017 1.001     3391     3162
CCSplenium_mothers_age        0.002     0.001   -0.000    0.004 0.999     5861     3128
CCSplenium_f1:HCC            -0.004     0.002   -0.007   -0.001 1.001     2689     2900
```

Box 2: Summary of the best model [M02a] coefficients. f1 = maternal prenatal depressive symptoms (e.g., EPDS factor score), HCC = maternal prenatal hair cortisol concentration; f1 × HCC = interaction between maternal prenatal depressive symptoms and maternal prenatal hair cortisol concentration; SEXtytto = female infants; KOKONAISIKA_VKO = infants post-menstrual age.

strong. Considering CC Body, the posterior is 88% within the ROPE — supporting the idea that the interaction is a bit stronger. Finally, considering CC Splenium, the posterior is 65% within the ROPE — highlighting that this interaction is the strongest. NOTE: Keep in mind that the choice of the ROPE was made subjectively.

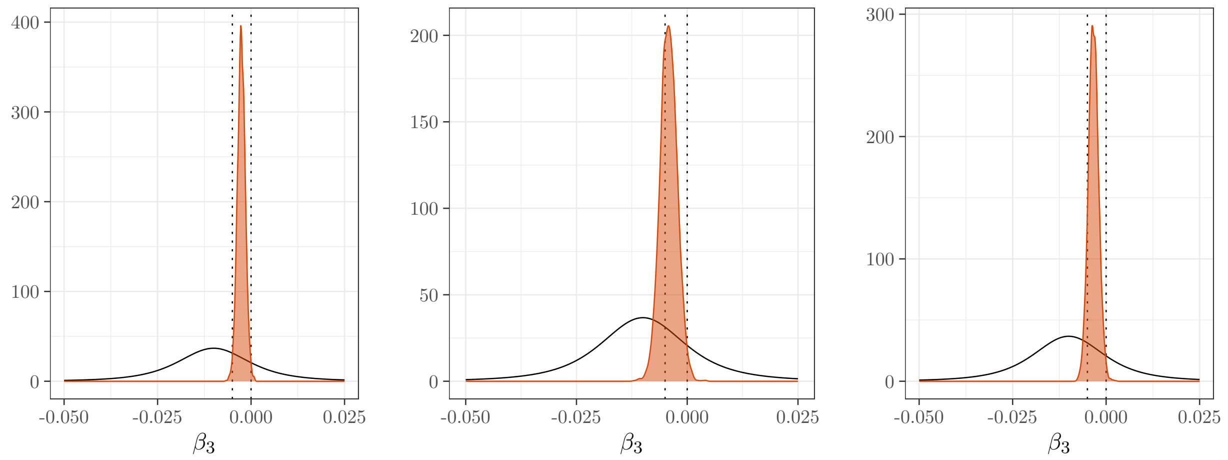

Figure S9: Interaction parameter: comparison between the prior (blue line) and the posterior (red shaded area). Dotted lines represent the ROPE bounds [-0.005, 0].

#### 4.5 Whole brain analysis

Figure S10 shown the results from the whole-brain regression model which were in line with our ROI results. However, the uncorrected T scores indicate that the effect is located in middle and anterior parts of the corpus callosum but only a small cluster was statistically significant after TFCE correction for the interaction model testing. The statistical map has been overlaid on the FA skeleton. [A] Only voxels reaching statistical significance ($p < 0.05$) are displayed, with threshold-free cluster enhancement corrected for multiple comparisons. [B] T map showing the T score for the same comparison. All figures are presented in neurological convention (left is left and right is right).

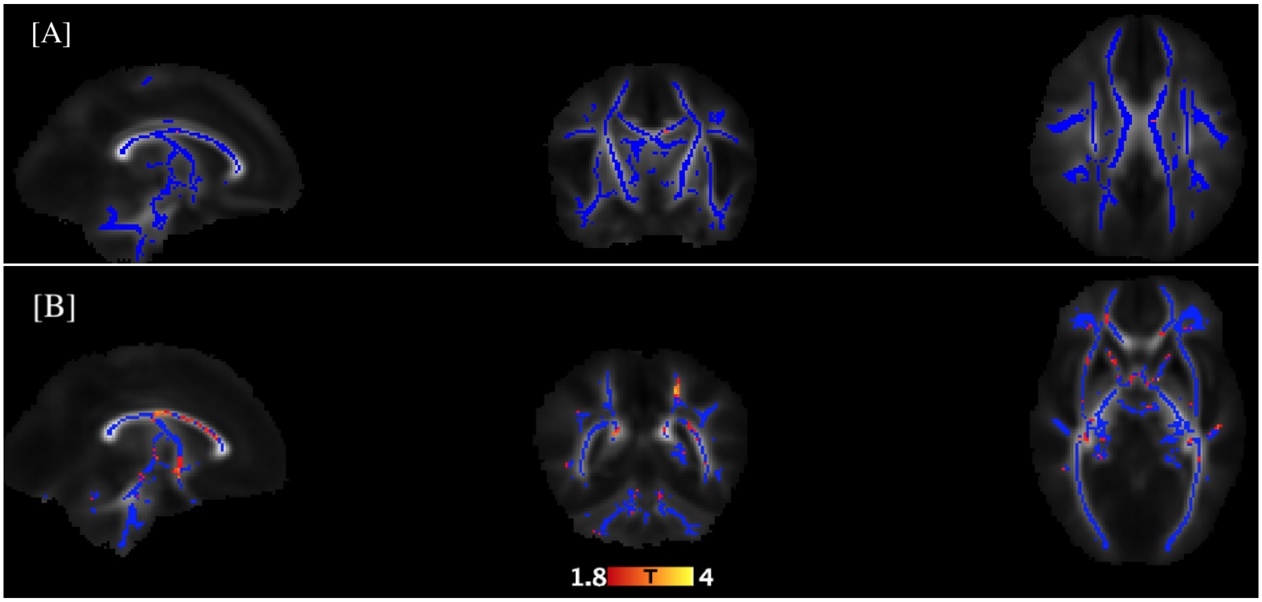

Figure S10: Whole brain analysis. [A] TFCE corrected result; [B] T map showing the T score for the same comparison.

### R packages

- brms. Bürkner, P. C. (2017). brms: An R package for Bayesian multilevel models using Stan. Journal of Statistical Software, 80, 1– 28.

- DataExplorer. Cui B (2020). DataExplorer: Automate Data Exploration and Treatment .R package version 0.8.2.

- ggdist. Kay M (2022). ggdist: Visualizations of Distributions and Uncertainty . doi: 10.5281/zenodo.3879620

- ggplot2. Wickham H (2016). ggplot2: Elegant Graphics for Data Analysis. Springer-Verlag New York. ISBN 978-3-319-24277-4.

- knitr. Xie Y (2022). knitr: A General-Purpose Package for Dynamic Report Generation in R . R package version 1.40

- psych. Revelle W (2023). psych: Procedures for Psychological, Psychometric, and Personality Research. Northwestern University, Evanston, Illinois. R package version 2.3.9.
